# Parallel evolution of tobramycin resistance across species and environments

**DOI:** 10.1101/758979

**Authors:** Michelle R. Scribner, Alfonso Santos-Lopez, Christopher W. Marshall, Christopher Deitrick, Vaughn S. Cooper

**Author notes:** Corresponding author: Vaughn Cooper.

## Abstract

An important problem in evolution is identifying the genetic basis of how different species adapt to similar environments. Understanding how various bacterial pathogens evolve in response to antimicrobial treatment is a pressing example of this problem, where discovery of molecular parallelism could lead to clinically useful predictions. Evolution experiments with pathogens in environments containing antibiotics combined with periodic whole population genome sequencing can be used to characterize the evolutionary dynamics of the pathways to antimicrobial resistance. We separately propagated two clinically relevant Gram-negative pathogens, *Pseudomonas aeruginosa* and *Acinetobacter baumannii*, in increasing concentrations of tobramycin in two different environments each: planktonic and biofilm. Independent of the pathogen, populations adapted to tobramycin selection by parallel evolution of mutations in *fusA1*, encoding elongation factor G, and *ptsP*, encoding phosphoenolpyruvate phosphotransferase. As neither gene is a direct target of this aminoglycoside, both are relatively novel and underreported causes of resistance. Additionally, both species acquired antibiotic-associated mutations that were more prevalent in the biofilm lifestyle than planktonic, in electron transport chain components in *A. baumannii* and LPS biosynthesis enzymes in *P. aeruginosa* populations. Using existing databases, we discovered both *fusA1* and *ptsP* mutations to be prevalent in antibiotic resistant clinical isolates. Additionally, we report site-specific parallelism of *fusA1* mutations that extend across several bacterial phyla. This study suggests that strong selective pressures such as antibiotic treatment may result in high levels of predictability in molecular targets of evolution despite differences between organisms’ genetic background and environment.

## Introduction

The notion that evolution can be forecasted at the level of phenotype, gene, or even amino acid is no longer a fantasy in the post-genomic era (Lässig et al., 2017). If we acknowledge that most forecasting efforts rely on history to anticipate the future, the explosive growth of whole-genome sequencing (WGS) sets the stage to resolve evolutionary phenomena in action and suggest the next selected path. Among the best examples, bacterial populations exposed to strong selection like antibiotics and analyzed by WGS are likely to identify gene regions that produce resistance (Ahmed et al., 2018a; Cooper, 2018; Feng et al., 2016; Palmer and Kishony, 2013). Repeated instances of the same antibiotic selection may enrich the same types of mutations and ultimately enable some measure of predictability (Ibacache-Quiroga et al., 2018; Wong et al., 2012). For instance, we can be confident that exposure of many bacteria to high doses of fluoroquinolones like ciprofloxacin may select for substitutions in residues 83 or 87 of the drug target, DNA gyrase A (Fàbrega et al., 2009; Wong and Kassen, 2011). Furthermore, effective prediction of drug resistance phenotypes based on genome sequence data has been demonstrated for certain bacterial species (Bradley et al., 2015; Tamma et al., 2019). These predictable outcomes are the product of very strong selection in populations with ample mutation supply and relatively few single mutations that can achieve high-level resistance (Ibacache-Quiroga et al., 2018).

Yet predicting evolution may be hampered when antibiotic selection produces species or environment-specific outcomes. Evolution experiments in antibiotics have demonstrated that subjecting different bacterial strains to the same antibiotic treatment regime (Gifford et al., 2018; Vogwill et al., 2014, 2016), or the same strains to different environments (Ahmed et al., 2018a; Santos-Lopez et al., 2019) can select for different drug resistance levels as well as molecular targets. We can test the extent of predictability of evolved levels and causes of drug resistance by studying the evolution of resistance in different environments and across different species that inherently have different genetic backgrounds. Here, we experimentally evolved two bacterial species from different genera, *Acinetobacter baumannii* and *Pseudomonas aeruginos*a, in increasing concentrations of the aminoglycoside tobramycin (TOB) in both planktonic and biofilm environments. We performed whole population genome sequencing throughout the history of each replicate lineage to determine their population-genetic dynamics and identify the molecular targets under selection in each condition.

*A. baumannii* and *P. aeruginosa* are ESKAPE pathogens that are responsible for multidrug-resistant infections (Santajit and Indrawattana, 2016). These species are members of the Moraxellaceae and Pseudomonadeae families and the strains used differ in their genome sizes by 2.7Mb, or more than 40%. Infections with these two opportunistic pathogens are often associated with a biofilm mode of growth (Eze et al., 2018; Mulcahy et al., 2014), where the bacteria grow in aggregates on surfaces and are protected from antimicrobials by a number of mechanisms. This biofilm protection may occur from secreted substances like polysaccharides, proteins, or eDNA that limit diffusion or by slowing growth and rendering the bacteria less susceptible to an antibiotic (Hall and Mah, 2017; Høiby et al., 2010). Given the lifestyle differences between cells growing in a biofilm compared to free-living cells, we asked whether evolution of TOB resistance could proceed by different mechanisms between these two environments. TOB is commonly used to treat infections caused by Gram-negative pathogens, and as an aminoglycoside can kill bacteria by multiple mechanisms (Bulitta et al., 2015). Aminoglycosides are actively transported into the cell following binding to the outer membrane, and subsequently can cause cell death by binding the 30S ribosome and blocking translation (Kohanski et al., 2010). They have also been shown to kill by binding to the outer membrane (Bulitta et al., 2015), and induce killing of cells that are not actively dividing (McCall et al., 2019).

In addition to identifying the range of molecular mechanisms of resistance available in these species and environments, we aimed to examine the evolutionary dynamics of adaptation in the presence of TOB. The success of the available molecular mechanisms of resistance is determined by the order in which causative mutations occurred (Wistrand-Yuen et al., 2018), the fitness imposed by those mechanisms (MacLean and Buckling, 2009), and the combinations of these mutations that are selectively tolerated (Knopp and Andersson, 2018). We used whole-population genome sequencing at regular intervals as we increased antibiotic concentrations to determine how selected mutations interact with the external environment and with one another, either competing on different haplotypes or combining in the same genotype.

## Results

We used TOB-sensitive ancestral clones of *A. baumannii* ATCC 17978 and *P. aeruginosa* strain PA14 to inoculate five replicate, single-species lineages for each of four treatments: planktonic without drug, planktonic with drug, biofilm without drug, and biofilm with drug. We propagated populations for twelve days and periodically froze samples for later sequencing and phenotypic analysis. The experimental design is illustrated in Figure 1A and 1B.

**Figure 1.**
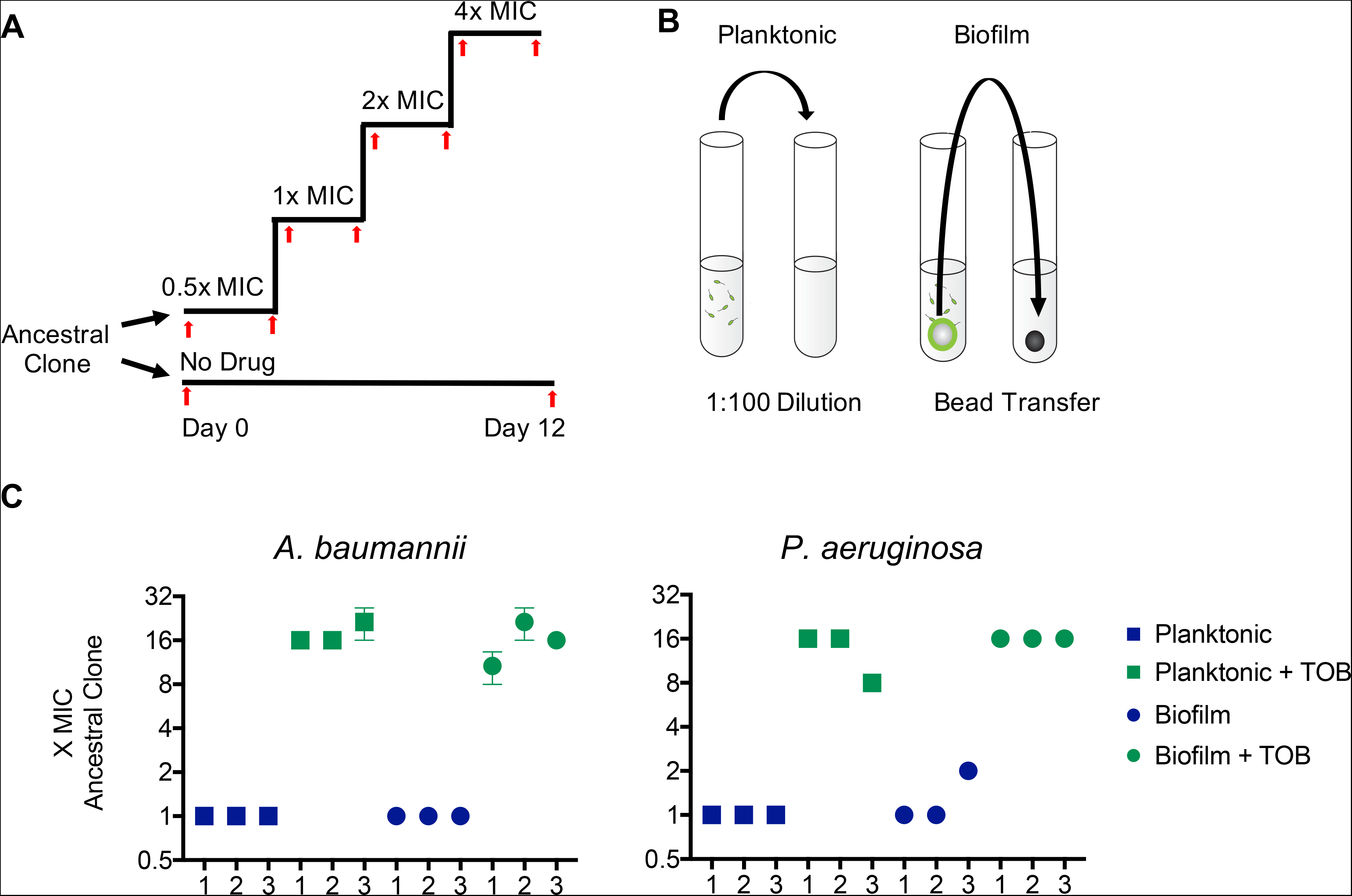
Parallel evolution of tobramycin resistance level across species and environment. Populations of *A. baumannii* and *P. aeruginosa* were propagated in minimal media with either increasing concentrations of tobramycin or no drug and either planktonic or biofilm lifestyle. Five replicate populations were propagated per treatment. **(A)** Populations were either propagated for twelve days in no antibiotic or inoculated into half the minimum inhibitory concentration of tobramycin with doubling concentrations every 72 hours. Samples of each population were archived for later phenotypic analysis and sequencing periodically throughout the experiment (red arrows). **(B)** Populations were propagated with either selection for planktonic growth through a daily 1:100 dilution or biofilm growth through a daily bead transfer that forces cells to undergo the entire biofilm lifecycle of attachment, growth, dispersion and reattachment every 24 hours as described in previous work (Poltak and Cooper, 2011). **C).** Tobramycin resistance level relative to the ancestral clone for three randomly chosen populations per treatment after twelve days of evolution. MICs were determined by microdilution in Mueller Hinton Broth according to CLSI guidelines. The mean of at least three replicate MIC assays per population is shown and error bars represent SEM. *A. baumannii* ancestral MIC = 1.0 mg/L, *P. aeruginosa* ancestral MIC = 0.5 mg/L in Mueller Hinton Broth. Populations must acquire resistance to 4x the MIC of the ancestral strain in order to survive the experiment (gray dashed line).

### Parallel evolution of TOB resistance phenotypes and genotypes

For a population to survive to the end of the experiment, it must evolve resistance to at least four times (4x) the TOB concentration that would kill the ancestral clone. Previous evolution experiments in antibiotics show that replicate populations may acquire different levels of resistance in response to the same antibiotic treatment regime when evolving in different environments (Gifford et al., 2018; Santos-Lopez et al., 2019; Trampari et al., 2019). While resistance levels did not change during the experiment for populations not exposed to antibiotics, populations that evolved with antibiotic selection survived TOB concentrations 8-21.3x MIC of the ancestral clone (Figure 1C and 1D).

Screens of transposon mutant libraries have identified 135 genes associated with low level resistance to aminoglycosides, suggesting that TOB resistance might arise by mutations in various molecular targets (Schurek et al., 2008). Instead, whole genome sequencing of the twenty TOB-treated populations at day twelve revealed that mutations in only a few loci rose to high frequencies (Figure 2, Figure S2). The large effective population sizes (>10^7^) of these experiments ensure that mutations occurred in nearly every position across the genome, and often multiple times (Santos-Lopez et al. 2019, Cooper, 2018). Therefore, the mutations identified by population-wide WGS, which in our case detect only those ≥5% frequency, represent the fittest resistance mutations of many contenders. The finding that mutations in the same genes evolved in parallel across antibiotic-treated populations (Figure 2) provides clear evidence of strong selection for their fitness benefits in the presence of TOB, and their absence in drug-free populations indicates they were not simply selected by other experimental conditions (Figure S2). In the unlikely possibility that these particular loci experienced significantly higher mutation rates in the presence of TOB, only selection would have driven them to these frequencies within 3-12 days (Cooper, 2018).

**Figure 2.**
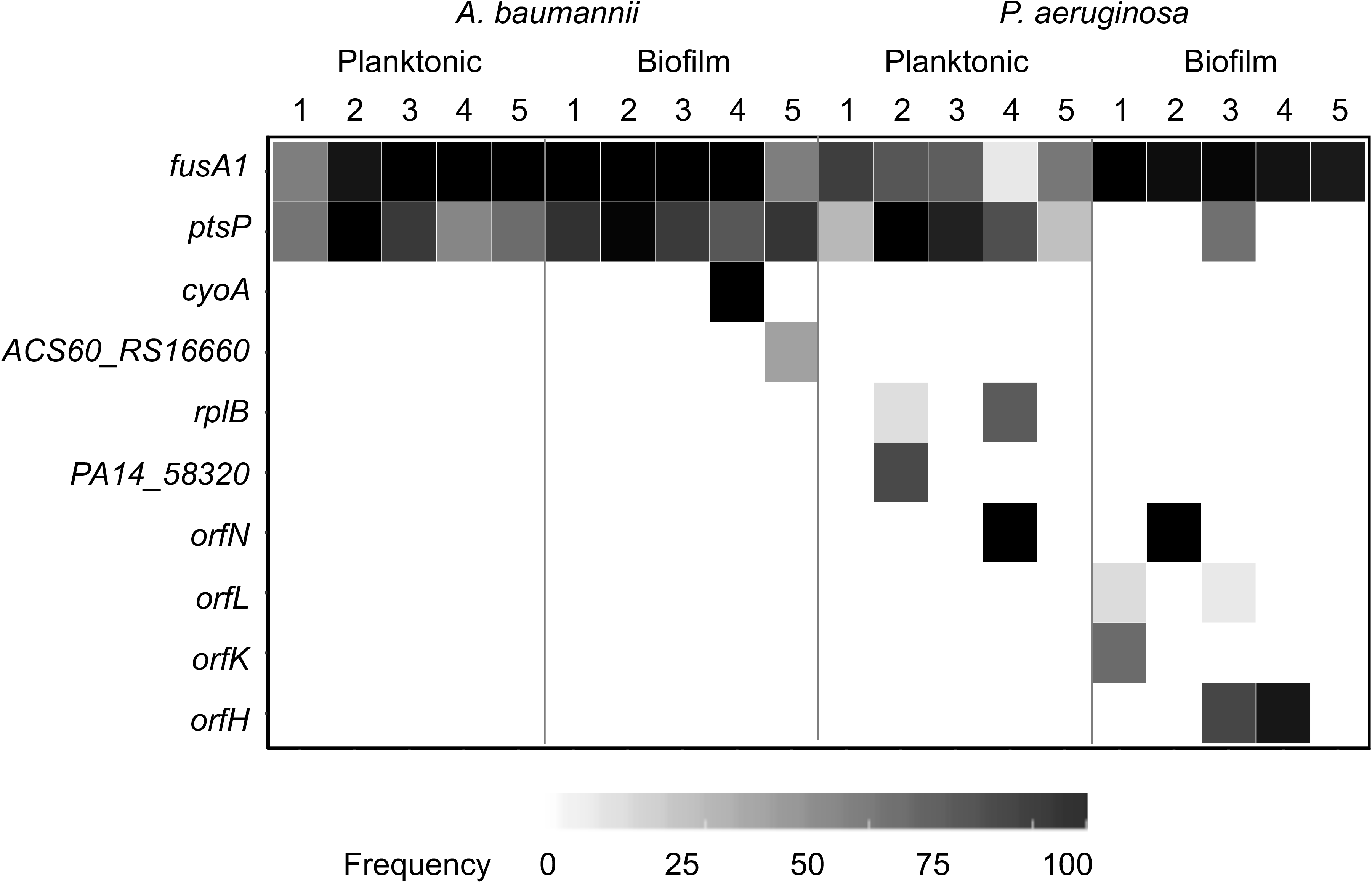
Population sequencing reveals interspecies parallelism and lifestyle-dependence of molecular targets of evolution. Tobramycin-associated mutations identified by whole population genome sequencing of *A. baumannii* and *P. aeruginosa.* Five populations per treatment were sequenced after 12 days of experimental evolution. Shading indicates the total frequency of all mutations in each gene within a population at day twelve.

Despite the many differences between *A. baumannii* and *P. aeruginosa*, both species frequently acquired mutations in *fusA1* and *ptsP* (Figure 2). The *fusA1* gene encodes elongation factor G (EF-G), an essential protein which functions in catalyzing translocation and ribosome recycling during translation (Savelsbergh et al., 2009). While A. *baumannii* has one copy of *fusA1*, *P. aeruginosa* and other *Pseudomonas* species also harbor an additional copy encoded by the gene *fusA2* (Palmer et al., 2013). EF-G is not known to be a direct binding target of TOB, and has received little attention as a mechanism of resistance to aminoglycosides in these species (Bolard et al., 2017; Sanz-García et al., 2018). However, this protein is the direct binding target of other antibiotics including fusidic acid and argyrin B (Johanson and Hughes, 1994; Jones et al., 2017). The exact mechanism by which mutations in EF-G confer aminoglycoside resistance is currently unknown but this study demonstrates it is an important resistance determinant. The *ptsP* gene encodes phosphoenolpyruvate phosphotransferase protein, which is part of the nitrogen phosphotransferase system and has previously been identified as a target of aminoglycoside resistance for *P. aeruginosa*, but not *A. baumannii* (Sanz-García et al., 2018; Schurek et al., 2008). The mechanism by which mutations in *ptsP* may confer resistance to TOB is also unknown, although the nitrogen phosphotransferase system has been shown to regulate expression of genes responsible for coordinating the antibiotic stress response, suggesting an indirect link between the nitrogen phosphotransferase system and antibiotic resistance (Gebhardt and Shuman, 2017). Therefore, despite as many as 135 genes in which mutations confer reduced susceptibility, mutations in these two genes *fusA1* and *ptsP* appear to be most fit in the presence of TOB across species and environments.

To clarify the specific effects of these mutations in TOB resistance, isogenic mutants were obtained by isolating clones from evolved *P. aeruginosa* populations and genotyping them by WGS. Two isogenic *fusA1* clones (N592I, Q678L), and *ptsP* clones (Δ14bp,1296-1309, Δ42bp, 1846-1887) were isolated and their resistance profiles measured. All were an average fourfold more resistant to TOB than the ancestral clone (Figure S3A). We also tested if mutations in *fusA1* increased the MIC to other ribosome-targeting antibiotics. These mutants were 2-4x more resistant to including amikacin, gentamycin, and tigecycline than the ancestor. Mutations in *ptsP* did not produce cross resistance to other antibiotics tested (Figure S3B), suggesting specificity to TOB resistance. While species-specific mutations did occur in *cyoA, cyoB*, *orfK, orfH, orfL, and orfN* as discussed below, the parallel evolution of mutations in *fusA1* and *ptsP* across *A. baumannii* and *P. aeruginosa* indicate that regardless of genetic background, these are among relatively few loci in which mutations can jointly increase TOB resistance and fitness in these conditions.

### Environment-associated adaptations to TOB selection

Experimental evolution in both planktonic and biofilm conditions allows us to test if the genetic pathways of adaptation depend on the external environment. TOB selection enriched multiple mutations in *fusA1* within each population regardless of the environment, but their frequencies differed with lifestyle. For *P. aeruginosa, fusA1* mutations dominated biofilm populations in the final sample (95.4%±3.7) but their frequencies varied in planktonic populations (50.4%±25.7) (Figure 2). Mutations in *ptsP* were prevalent in planktonic populations but only one biofilm population contained a *ptsP* mutation at day twelve. Rather, these biofilm populations of *P. aeruginosa* biofilm populations frequently acquired mutations in *orfK, orfH, orfL*, or *orfN* genes (subsequently referred to jointly as *orfKHLN*), encoding O antigen biosynthesis enzymes, whereas this locus was only mutated in one of the planktonic populations (Burrows et al., 1996; Rocchetta et al., 1999). A mutation in *orfK* also occurred in a population with biofilm but no TOB selection, suggesting that *orfKHLN* mutations may be beneficial in both biofilm and TOB selection alone, but most beneficial in the combination of these conditions (Figure 2).

The genetic targets of resistance were more consistent in *A. baumannii* populations treated with TOB, with *fusA1* and *ptsP* mutations reaching similar frequencies in both biofilm and planktonic treatments by the end of the experiment. However, mutations in *cyoA* and *cyoB* (subsequently referred to jointly as *cyoAB*), encoding components of the electron transport chain, were associated with only biofilm lineages (Figure 2) (Ibacache-Quiroga et al., 2018). Together, the mutations enriched in biofilm lineages (*orfKHLN, cyoAB*) indicate that lifestyle may influence the identity of targets under selection by TOB or their frequency (*fusA1, ptsP*).

The parallel evolution of mutations in four genes associated with O antigen biosynthesis within and between populations strongly suggests that the outer membrane may be altered in response to TOB and biofilm selection in *P. aeruginosa* (Burrows et al., 1996; Tognon et al., 2017; Wong et al., 2012). Experiments in strain PAO1 have shown that loss of B band O antigen is associated with aminoglycoside resistance by increasing impermeability and reducing binding affinity to the outer membrane (Bryan et al., 1984; Kadurugamuwa et al., 1993). Similarly, populations of *A. baumannii* evolved with TOB and biofilm selection acquired mutations in the *cyoAB* operon, encoding components of the electron transport chain. Mutations in electron transport chain components have previously been associated with resistance to aminoglycosides through increased membrane impermeability (Bryan and Kwan, 1983; Damper and Epstein, 1981; Schurek et al., 2008). Therefore, although the biofilm-associated mutations in the two species propagated in this experiment evolved mutations in different genetic loci (affecting LPS biosynthesis genes in *P. aeruginosa* and electron transport chain components in *A. baumannii*), these may represent parallelism in a broad strategy of altered membrane structure or permeability that is most beneficial under both biofilm and aminoglycoside selection.

### Population-genetic dynamics of TOB resistance evolution

We used longitudinal population sequencing of three lineages per treatment to determine effects of species and environment on the population-genetic dynamics of adaptation to the antibiotic. The trajectories of allele frequencies (shown in Figure S4) were used to predict mutation linkage and hence the assembly and dynamics of genotypes, which are represented by Muller plots (Figure 3, see methods). Genotype frequency is represented by the breadth of shading with colors corresponding to the putative driver mutations of that genotype. In all lineages, regardless of environment, mutations in *fusA1* were detected at either 0.5x MIC or 1.0x MIC and subsequently rose to high frequencies (Figure 3, red). Their rapid rise in frequency in the first few days of the experiment suggests that *fusA1* mutations were the fittest contending mutations at subinhibitory concentrations of TOB. Lineages with different SNPs in the *fusA1* gene coexisted in some populations for the duration of the experiment, suggesting that different *fusA1* genotypes had similar fitness in increasing TOB. While a single, selected nonsynonymous *fusA1* mutation was presumably sufficient for survival to the end of the experiment due to the presence of clones with no other mutations at day twelve (Supplementary Data), secondary mutations were selected in these genotypes in the genes discussed previously (*ptsP*, *orfKHLN*, and *cyoAB*, Figure 3). *A. baumannii* populations tended to become dominated by *fusA1+ptsP* genotypes, but biofilm populations also selected *cyoAB* mutants prior to day nine (up to 2x MIC) that were outcompeted by a *fusA1+ptsP* genotype at 4x MIC TOB. Planktonic populations of *P. aeruginosa* demonstrated coexistence of *fusA1*, *ptsP*, and *fusA1+ptsP* haplotypes throughout the experiment, rather than a sweep of a *fusA1+ptsP* genotype (Figure 3). In contrast, *P. aeruginosa* biofilm populations repeatedly selected *orfKHLN* mutants on a *fusA1* background. These evolutionary dynamics demonstrate that following initial selection of a *fusA1* mutation, selection favored secondary mutations particular to lifestyle and species.

**Figure 3.**
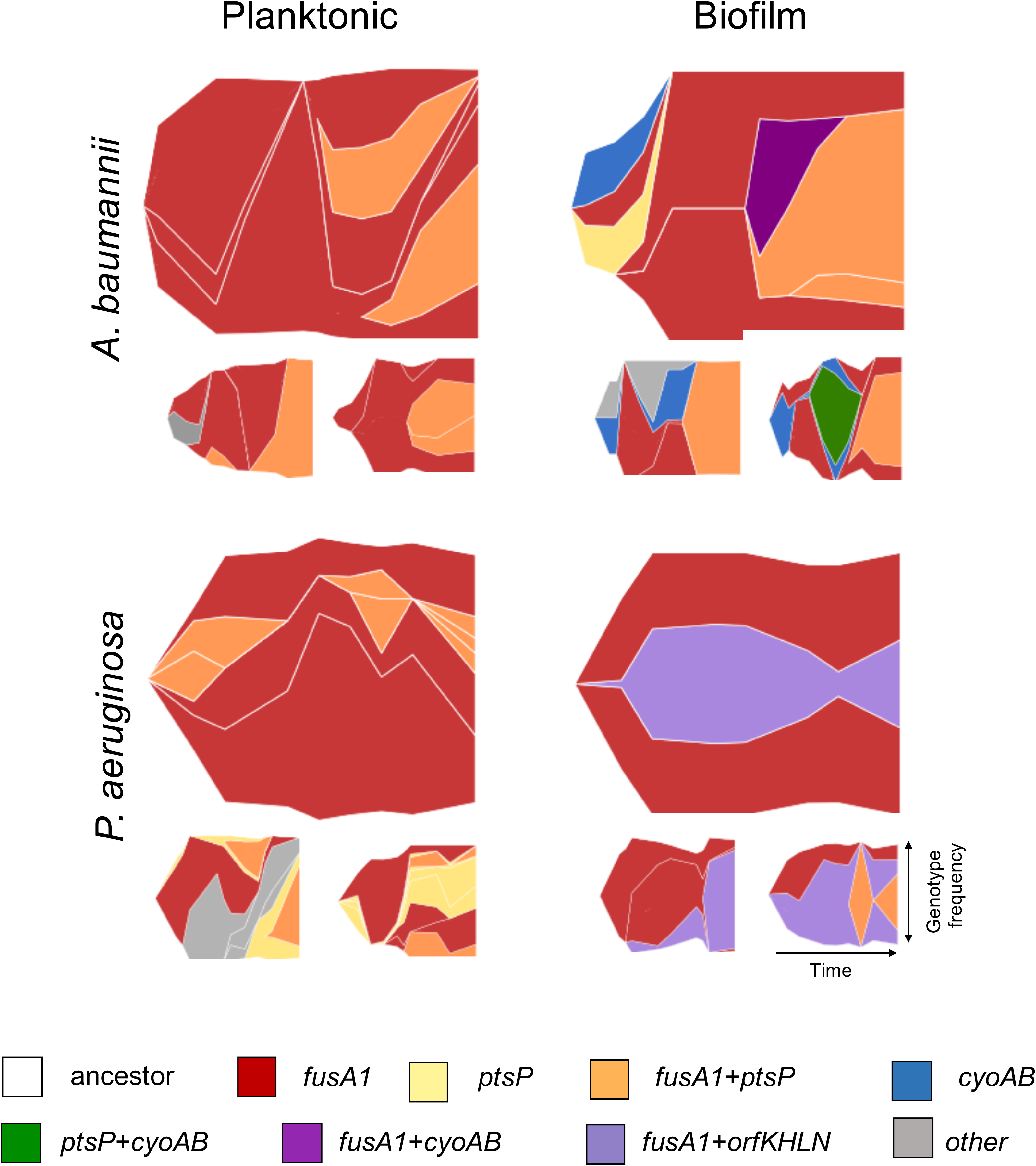
Evolutionary dynamics of bacterial populations in increasing concentrations of tobramycin. Muller diagrams displaying genotype frequencies as a proportion of the population throughout twelve days of evolution for three populations per treatment. Genotypes are shaded by the putative driver loci that are mutated. Different lineages of the same color represent mutations at different positions within the same loci that are coexisting within the population. The frequency of genotypes at every time point is represented by the height of the graph that it spans at that time point. In situations where a first mutation arises in the background of the ancestral genotype, the color representing that genotype can be seen beginning from the white background, whereas in situations where a mutation arises in the background of another mutation, thus generating a new genotype, the new color arises in the middle of the existing genotype. Mutations occurring in the background of putative driver mutations are not shown but may be viewed in linear allele frequency plots of each population in Figure S4.

### Parallelism of aminoglycoside resistance mechanisms across species and clinical isolates

The repeated evolution of *fusA1* and *ptsP* mutations in both *P. aeruginosa* and *A. baumannii* suggested that these mutations may provide a general mechanism of TOB resistance across diverse species. We tested this hypothesis by searching published datasets and genomes for *fusA1, ptsP, cyoA*, and *cyoB* mutations (Methods). Mutations in *fusA1* were found in several different species including *E. coli*, *S. typhimurium*, and *S. aureus* (Ibacache-Quiroga et al., 2018; Jahn et al., 2017; Johanson and Hughes, 1994; Kim et al., 2014; Mogre et al., 2014; Norström et al., 2007), and all laboratory studies reported these mutations either in response to aminoglycoside selection or as a direct cause of aminoglycoside resistance (Figure 4). Mutations in *cyoA* and *cyoB* were also found in *E. coli* and *S. typhimurium* in these experiments (Ibacache-Quiroga et al., 2018; Jahn et al., 2017; Johanson and Hughes, 1994). Multiple sequence alignment of these genes across species revealed that *fusA1* mutations were localized to two primary regions across all species and were primarily nonsynonymous SNPs, and eight positions exhibit amino acid-level parallelism across species (Figure 4, Table S1, Figure S5) (Wattam et al., 2017). In this dataset, mutations occurred at positions with identical amino acids across species more frequently than expected by chance for *fusA1* (χ^2^ = 11.58, df = 1, p = 0.0006), *ptsP* (χ^2^ = 4.37, df = 1, p = 0.03652), and *cyoA* (χ^2^ = 3.89, df = 1, p = 0.0486). The parallel evolution of *fusA1* mutations across species and environments demonstrates the potential utility of this gene as a predictive marker of aminoglycoside resistance.

**Figure 4.**
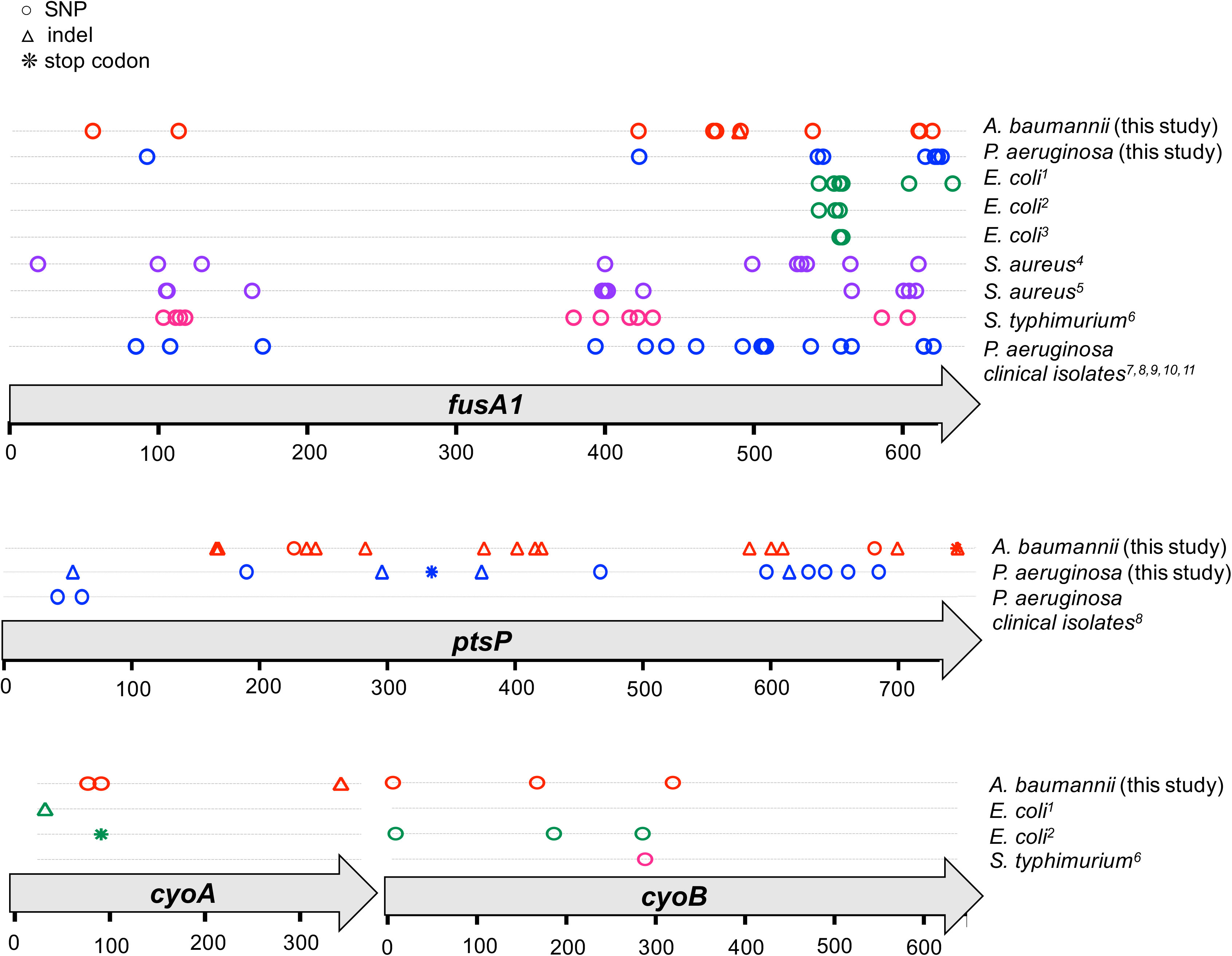
Parallelism of mutations in genetic loci associated with aminoglycoside resistance across species and clinical isolates. All mutations that occurred at any point in the experiment within the *fusA1*, *ptsP*, *cyoA*, or *cyoB* genes are indicated by a symbol at its position within the encoded amino acid sequence. Mutations reported in previous literature in other species are indicated and color coded by species; these mutations were either selected by aminoglycoside treatment *in vitro* or selected by another antibiotic and subsequently demonstrated to confer resistance to aminoglycosides. Mutations reported in whole genome sequencing datasets of *P. aeruginosa* clinical isolates are indicated. SNPs are indicated by a circle, insertions or deletions (indels) are indicated by a triangle, and stop codon mutations are indicated by an asterisk. Top: *fusA1* gene, Middle: *ptsP* gene, Bottom: *cyoA* and *cyoB* genes. >Referenced literature: 1. Jahn et al., 2017, 2. Ibacache-Quiroga et al., 2018, 3. Mogre et al., 2014, 4. Kim et al., 2014, 5. Norström et al., 2007, 6. Johanson and Hughes, 1994, 7. López-Causapé et al., 2017, 8. Markussen et al., 2014, 9. Chung et al., 2012, 10. Bolard et al., 2017, 11. López-Causapé et al., 2018

To examine if the precise mutations found in our *in vitro* study were also found in clinical isolates, we searched published genomes of *P. aeruginosa* clinical isolates from cystic fibrosis patients who had likely been treated with aminoglycosides like TOB (Bolard et al., 2017; Chung et al., 2012; López-Causapé et al., 2017, 2018; Markussen et al., 2014). We identified *fusA1* and *ptsP* mutations in these genomes, suggesting that these mutations evolve during infections. Although it is not possible to distinguish aminoglycoside selection as the driver of these mutations in a clinical setting, the selection of mutations in these genes in our evolution experiment and the increased resistance that these mutations have the same effect *in vivo*. Taken together, the parallel evolution of mutations in these genes in clinical isolates of a wide range of species indicates they commonly contribute to aminoglycoside resistance in diverse environments, including the cystic fibrosis respiratory tract.

## Discussion

The rapidly intensifying problem of antimicrobial resistance demands understanding of how antibiotic resistance evolves and which types of mutations or mobile elements are common causes (Brockhurst et al., 2019). Genetic screens of mutant collections have revealed potential resistance mechanisms (Schurek et al., 2008), and more recently, evolve-and-resequence experiments have been used to identify the most fit resistance mutations in a given condition (Santos-Lopez et al., 2019; Sanz-García et al., 2018; Wong et al., 2012). However, the broader clinical utility of these screens for predicting the evolution of antibiotic resistance depends upon the relevance of the findings in other genetic backgrounds or environments. This study served the dual purpose of identifying mutations that contribute to TOB resistance in *A. baumannii* and *P. aeruginosa* and demonstrating effects of different environments and species history on evolutionary dynamics and causes of resistance.

Because antibiotics often induce cell killing by targeting conserved domains of proteins involved in essential cell processes, there is potential for mutations in these genes to confer resistance in diverse bacterial strains and species. However, many have appreciated that epistatic interactions stemming from differing genetic backgrounds could limit parallel evolution of resistance mechanisms (Breen et al., 2012; MacLean et al., 2010; Ward et al., 2009). In spite of the many genetic differences between *A. baumannii* and *P. aeruginosa* – the latter genome much larger and containing dozens of additional putative resistance loci – we identified parallel mutations in *fusA1* and *ptsP* in response to tobramycin selection, a largely unknown combination of mutations conferring high fitness and resistance. Furthermore, we found amino acid-level parallelism of *fusA1* mutations associated with aminoglycoside resistance, including kanamycin, gentamycin, and amikacin, across diverse species including *E. coli*, *S. typhimurium*, and *S. aureus* (Figure 4, Figure S5). Our finding that *fusA1* mutations repeatedly evolve in clinical *P. aeruginosa* isolates from different strain types and host conditions further supports the notion that these mutations can arise in a range of genetic backgrounds and environments, and we predict that *fusA1* mutations may be considerably more prevalent following antibiotic therapy than previously appreciated. These *fusA1* mutations also produced cross resistance to other ribosome-targeting antibiotics, a concerning finding given the frequent use of tobramycin in treating infections of the CF airway (Figure S3) (Chmiel et al., 2014).

Tobramycin selection in a biofilm model of growth demonstrably altered the targets of selection in ways that motivate studies of the mechanism of aminoglycoside killing in this lifestyle. The rise of mutations in LPS biosynthesis genes (*orfKHLN*) and electron transport chain components (*cyoAB*) primarily in biofilm populations indicates that altered binding or cell permeability may have been selected and can indicate a distinct mode of action of TOB. These lifestyle distinctions in resistance traits suggest that the environment may influence the evolutionary dynamics of antimicrobial resistance, in concordance with previous studies (Ahmed et al., 2018b; Santos-Lopez et al., 2019; Trampari et al., 2019). Therefore, while genes like *fusA1* represent mechanisms of resistance that are robust across a wide range of species, environments, and host conditions, the prevailing mode of bacterial growth is likely crucial in attempting to predict the evolution of antimicrobial resistance.

The extent of parallelism in molecular evolution in these experiments is surprising and its causes need to be considered. Why would widely different species evolve to resist tobramycin by *fusA1* mutations, and to a lesser extent *ptsP* mutations, when many causes of aminoglycoside resistance are likely available (Schurek et al., 2008)? Several possible explanations exist that are not mutually exclusive. One possibility is that these genes possess a high local mutation rate and thus acquire mutations more rapidly than other available molecular targets of resistance. However, it is doubtful that these mutations are more available than others, since neither were enriched in studies of mutations accumulated in the near-absence of selection nor are these loci in genome regions shown to have higher mutation rates (Dettman et al., 2016; Long et al., 2014). More likely is that mutations to *fusA1* and *ptsP* simply produce the greatest fitness benefit in these conditions and these fitness benefits are robust to different species and environments. Other drugs may have a wider range of targets that produce the same level of fitness benefit, resulting in less parallelism. The target size of these genes, in which multiple nonsynonymous mutations produce resistance, may also contribute to gene-level parallelism across populations: with a larger proportion of available beneficial mutations at these concentrations of tobramycin, gene-level parallelism in *fusA* and *ptsP* may be more likely.

Another cause of molecular parallelism is that when the fittest available molecular targets of resistance are in highly conserved genes, such as *fusA1* (Savelsbergh et al., 2009), mutations in these loci may be predictable across otherwise highly divergent species. Conversely, if a strain gains a highly fit resistance mechanism using genes that are not conserved, gene-level parallelism with other strains is not possible and the target predictability diminished. The exact mechanism by which alterations in elongation factor G produce resistance is not known, but have been suggested to produce structural changes that could interfere with aminoglycoside binding (Bolard et al., 2017). Nonetheless, the nucleotide-level parallelism we have observed suggest that the effect of these changes may be conserved across species and inform the molecular mechanism of resistance. The finding of *fusA1* mutations across species also suggests that these mutations may have relatively low epistatic interactions with the genetic background of the strain. Predictability may be more challenging if epistatic effects of resistance alleles are greater.

Several studies of experimentally evolved microbial populations have demonstrated a general “rule of declining adaptability”, whereby beneficial mutations produce diminishing fitness benefits as a population becomes better adapted to its environment (Kryazhimskiy et al., 2014; Wang et al., 2016). These studies have experimentally determined that populations exhibit global diminishing returns epistasis, in which the relative fitness level of different strains in an environment influences the fitness effect of a given mutation more than the specific genetic differences between the strains. Our study is consistent with this model: *A. baumannii* and *P. aeruginosa* are highly unfit in the TOB concentrations utilized in this study and exhibit similar benefits from *fusA1* and *ptsP* mutations despite genetic differences in the ancestral genotypes. Theory suggests that if these strains were more fit in these conditions, for example if they were more resistant to the TOB concentrations in which they are growing, *fusA1* and *ptsP* mutations may be less beneficial and thus evolution less predictable. In other antibiotic or species combinations, it remains to be seen if effects of resistance mutations are heavily skewed toward a few that are most fit and whether subsequent resistance mutations exhibit diminishing returns that weaken selection and increase variation among populations.

In this study we have identified gene and nucleotide level parallelism of molecular targets of tobramycin resistance across species, environments, and clinical isolates. While environmental and genetic differences are crucial factors influencing the evolutionary dynamics of resistance, this parallelism indicates that the strong selective pressures produced by antibiotics may result in strikingly predictable targets of molecular evolution in certain drugs and species combinations.

## Methods

### Strains and Media

*Pseudomonas aeruginosa* strain UCBPP-PA14 and *Acinetobacter baumannii* strain ATCC 17978 were the ancestral strains used in the evolution experiments (Baumann et al., 1968; Piechaud and Second, 1951; Rahme et al., 1995). *A. baumannii* ATCC 17978 was propagated for ten days in minimal media to pre-adapt it to the media conditions prior to the evolution experiment. The minimal media used in the evolution experiments consisted of an M9 salt base (0.37 mM CaCl_2_, 8.7 mM MgSO_4_, 42.2 mM Na_2_HPO_4_, 22 mM KH2PO_4_, 21.7mM NaCl, 18.7 mM NH_4_Cl), 0.2 g/L glucose, 20 mL/L MEM essential amino acid solution, 10 mL/L MEM nonessential amino acid solution (Thermofisher 11130051, 11140050), and 1 mL/L each of Trace Elements A, B, and C (Corning 99182CL, 99175CL, 99176CL). In addition, DL-lactate (Sigma-Aldrich 72-17-3) was added to the *P. aeruginosa* medium to a final concentration of 10mM in order to generate the approximate nutrient concentrations present in the cystic fibrosis lung environment (Palmer et al., 2007).

### Evolution Experiment

Evolution experiments in both *P. aeruginosa* and *A. baumannii* were initiated using a single ancestral clone. For *P. aeruginosa*, a single colony was selected and resuspended in PBS, then used to inoculate twenty replicate lineages. For *A. baumannii*, a single colony was selected and grown in minimal medium with no antibiotic for 24 hours, then used to inoculate twenty replicate lineages. Lineages were propagated with either increasing concentrations of TOB or no TOB and either planktonic or biofilm selection, such that five replicate lineages each were propagated for four experimental conditions (planktonic without TOB, planktonic with TOB, biofilm without TOB, and biofilm with TOB) for each organism. Lineages with planktonic selection were propagated through a 1:100 dilution every 24 hours (50uL into 5mL of fresh minimal media), and lineages with biofilm selection were propagated through a transferring a colonized polystyrene bead (Cospheric, Santa Barbara, CA) to a tube of fresh media and three fresh beads every 24 hours, as described previously (Poltak and Cooper, 2011). *P. aeruginosa* biofilm transfers were performed by transferring a bead directly to the next day’s tube, whereas *A. baumannii* biofilm transfers were performed by first rinsing the bead by transferring it to a tube of PBS, then to the next day’s tube. Lineages propagated with antibiotic selection were treated with tobramycin sulfate (Alfa Aesar, Wardhill, MA) starting at 0.5X MIC of the ancestral strain in the experimental minimal media (0.5 mg/L for A. *baumannii* and 2.0 mg/L for *P. aeruginosa*), with doubling of the concentration every 72 hours. The experiment was performed for twelve days, with samples collected on days 3, 4, 6, 7, 9, 10, and 12 and frozen at −80°C in either 25% glycerol for *P. aeruginosa* or 9% DMSO for *A. baumannii*. Planktonic lineages were sampled by freezing an aliquot of the liquid culture, and biofilm lineages by sonicating a bead in PBS and freezing an aliquot of the resuspended cells.

### Minimum Inhibitory Concentration Assays

We determined MICs by broth microdilution in Mueller Hinton Broth according to Clinical Laboratory Standards Institute guidelines (CLSI, 2019). To measure MIC’s for evolved populations, we revived frozen populations by streaking onto a ½ T-Soy agar plate, resuspended a portion of the resulting bacterial lawn in PBS, and diluted to a 0.5 McFarland standard. We inoculated the suspension into a round bottom 96 well plates containing two-fold dilutions of TOB at a final concentration of 5×10^5^ CFU/mL. *P. aeruginosa* and *A. baumannii* MIC assays were then incubated at 37°C for 16-20 hours or 18-22 hours, respectively, then the MIC was determined as the first well that showed no growth. At least three assays were performed for each population. In *P. aeruginosa*, MICs of TOB differed when measured in Muller Hinton Broth compared to the experimental minimal medium but reflect similar fold changes in MIC relative to the ancestral clone (Figure S1). Clones were measured by the same procedure with the exception that freezer stocks were streaked for isolation and MIC assays were performed using an isolated colony. MICs of other ribosome-targeting antibiotics for the *fusA1* and *ptsP* isogenic mutants were performed using Sensititre plates according to manufacturer specifications (Sensititre GN3F, Trek Diagnostics Inc., 514 Westlake, OH).

### Genome Sequencing and Analysis

Whole populations were sequenced periodically throughout the experiment. For TOB-treated lineages, all populations were sequenced on day 12, and 3 lineages from each of the planktonic and biofilm conditions were also sequenced on days 3, 4, 6, 7, 8, 9, and 10 for *P. aeruginosa* and days 1, 3, 4, 6, 7, 9, 10, and 12 for *A. baumannii*. Three no TOB lineages from each of the biofilm and planktonic conditions were also sequenced on days 6 and 12 for *P. aeruginosa* and days 1, 4, 9, and 12 for *A. baumannii*.

Populations were prepared for sequencing by inoculating freezer stocks of the bacterial populations into the same media and antibiotic concentration in which the population was growing in at the time of freezing. Identical growth conditions to the population’s growth conditions at the time of freezing were maintained in order to minimize bias in the population structure during the outgrowth process. After 24 hours of growth, populations were sampled by either removing an aliquot of the culture for planktonic populations or transferring beads to PBS, sonicating, and removing an aliquot of the resuspended cells for biofilm populations. DNA was extracted using the DNeasy Blood and Tissue Kit (Qiagen, Hiden, Germany). The sequencing library was prepared as described by Turner and colleagues (Turner, Marshall et al. 2018) according to the a previously described protocol (Baym et al., 2015) using the Illumina Nextera kit (Illumina Inc., San Diego, CA) and sequenced using an Illumina NextSeq500. Samples were sequenced to 160x coverage on average for *P. aeruginosa* populations and 309x coverage on average for *A. baumannii* populations.

Sequences were trimmed using the Trimmomatic software v0.36 (Bolger, Lohse et al. 2014) with the following criteria: LEADING:20 TRAILING:20 SLIDINGWINDOW:4:20 MINLEN:70. The breseq software v0.31.0 was used to call variants using the default parameters and the -p flag when analyzing population sequences. These parameters call mutations only if they are present at least 5% frequency within the population and are in at least 2 reads from each strand. The *A. baumannii* ATCC 17978-mff genome (NZ_CP012004) and plasmid NZ_CP012005 sequences were downloaded from RefSeq. Two additional plasmids were found to exist in our working strain and were added to this reference genome: NC009083 and NC_009084. The *P. aeruginosa* UCBPP-PA14 genome was downloaded from RefSeq (NC_008463). Mutations were removed if they were also found in the ancestor’s sequence when mapped to the reference genome. Mutations that did not reach at least 25% cumulative frequency across all populations at all timepoints were removed, and mutations were also manually curated to remove biologically implausible mutations. A mutation was determined biologically implausible if it occurred either *i*) at trajectories that were not possible given the trajectories of the putative driver mutations, *ii*) at only the ends of reads, only reads with many other mutations, or at only low coverage (<10 reads), indicating poor read mapping at that region. When high-quality mutations in loci related to the putative driver loci or ribosome machinery were reported in New Junction Evidence by breseq, these mutations were also included in the analysis. Mutations fitting these criteria included mutations to 23S rRNA in *P. aeruginosa*, and mutations to *ptsP*, HPr, and NADH quinone oxidoreductase in *A. baumannii*. Filtering, allele frequencies, and plotting were done in R v3.5.3 (www.r-project.org) with the packages ggplot2 v2.2.1 (https://CRAN.R-project.org/package=ggplot2) and tidyr (https://CRAN.R-project.org/package=tidyr). Muller plots were generated using the muller_diagrams package by CD (https://github.com/cdeitrick/muller_diagrams) v0.5.2 using default parameters. These scripts predict genotypes and lineages based on the trajectories of mutations over time using a hierarchical clustering method and implement filtering criteria to eliminate singletons that do not comprise prevalent genotypes. Muller plots were manually color coded by the presence of putative driver mutations within each genotype. Additional mutations that occurred on the background of putative driver mutations can be viewed in the allele frequency plots but were not shown in Muller plots (Figure 3, Figure S3).

### Resistance Loci Alignment and Mutation Mapping

Mutations in putative resistance loci (*fusA1*, *ptsP*, *cyoA*, and *cyoB)* were identified in previous literature reporting that these mutations arose in response to aminoglycoside selection or directly conferred an increase in aminoglycoside resistance by MIC assay. Amino acid sequences of the encoded proteins for these species were obtained by searching the “Features” section of PATRIC for these genes in the genomes specified in these experiments (Wattam et al., 2017). The amino acid sequences were aligned in PATRIC and mutations reported in each study were mapped to the corresponding position in the sequence alignment (Figure 4). Mutations in these genes in clinical isolates were found by searching for whole genome sequencing studies of *P. aeruginosa* isolates from cystic fibrosis patients that reported these mutations. We tested whether the SNPs identified in each of these genes occur in conserved positions more frequently than expected considering the frequency of conserved positions within the genes using a Pearson’s Chi-squared test (Figure S5).

## Acknowledgments and Funding Sources

We thank Daniel Snyder and the Microbial Genome Sequencing center (MiGS) for technical support. This work was supported by the Institute of Allergy and Infectious Diseases at the National Institutes of Health (grant U01AI124302 and grant T32AI049820) and by the Cystic Fibrosis Foundation Research Development Program.

