## Supplemental Figures and Tables for "Parallel evolution of tobramycin resistance across species and environments"

**Supplementary Material**

**Figures S1-S5**

**Table S1**

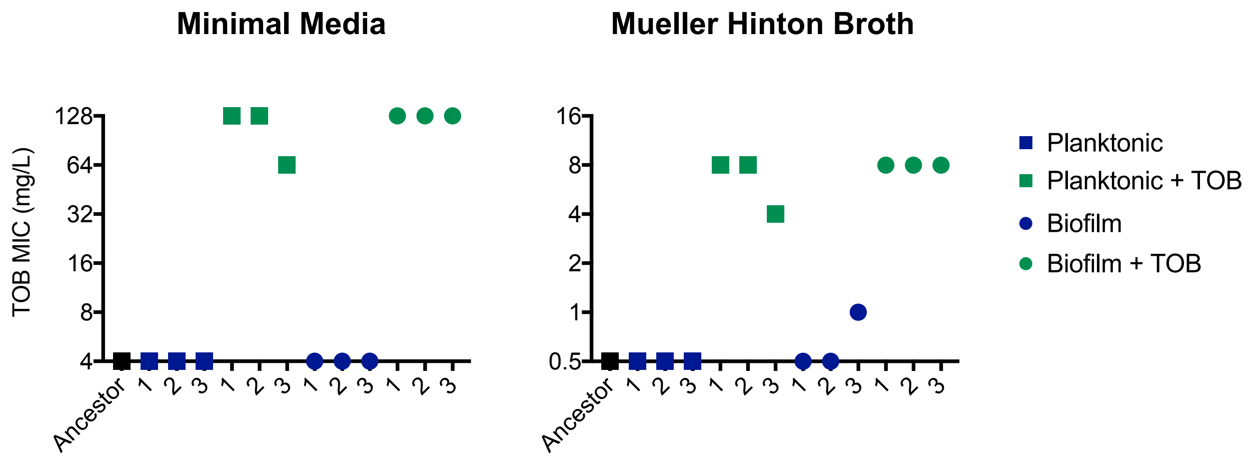


**Figure S1. Minimum inhibitory concentration of *P. aeruginosa* evolved populations on day twelve in the minimal media used in the evolution experiment (Left) and Mueller Hinton Broth (Right).** MICs were performed according to CLSI guidelines with the exception of media used. The mean of at least three replicate MIC assays per population is shown and error bars represent SEM.

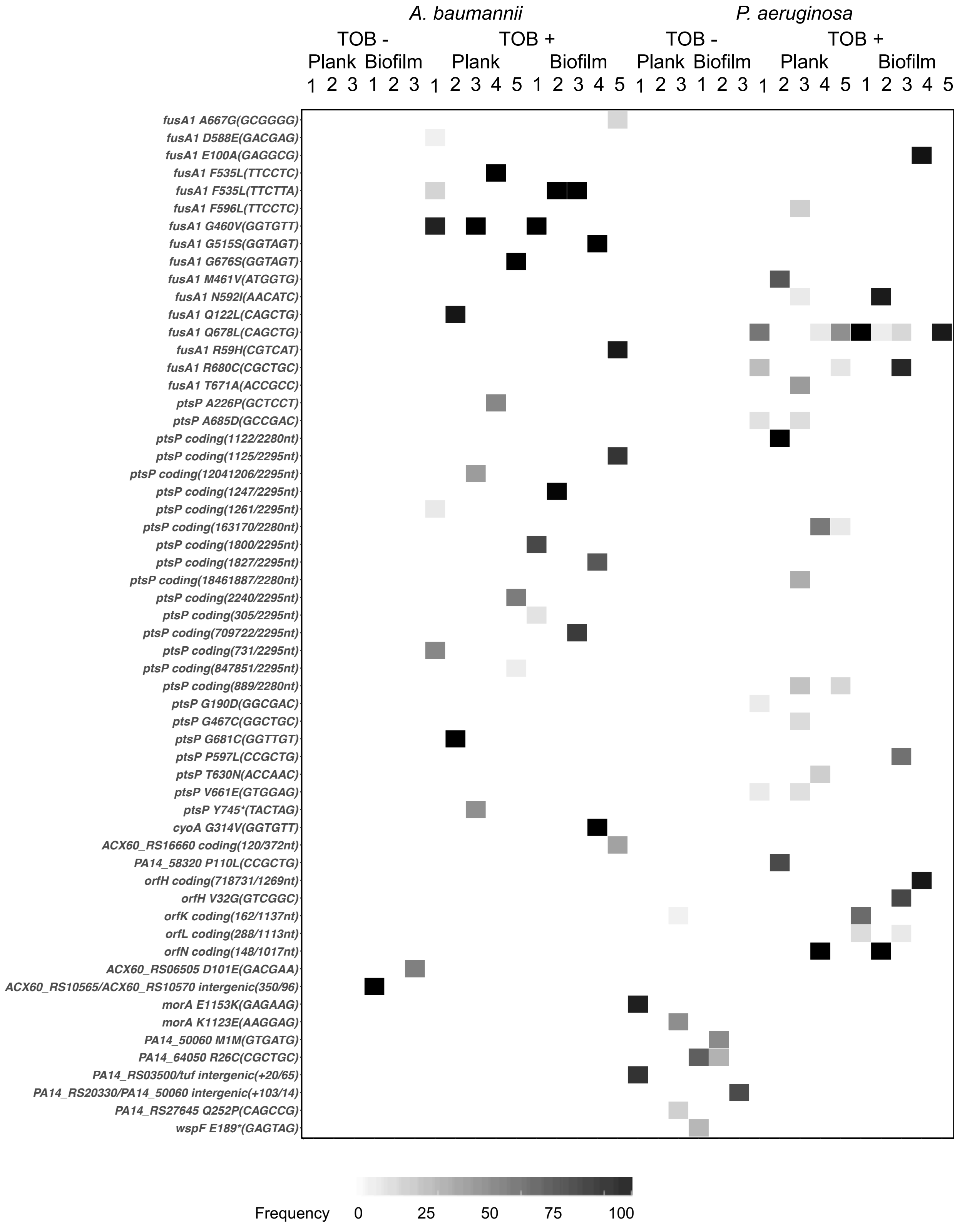
A. baumannii
P. aeruginosa
TOB -
TOB +
TOB -
TOB +
Plank
Biofilm
Plank
Biofilm
Plank
Biofilm
Plank
Biofilm
fusA1 A667G(GCGGGG)
fusA1 D588E(GACGAG)
fusA1 E100A(GAGGCG)
fusA1 F535L(TTCCTC)
fusA1 F535L(TTCTTA)
fusA1 F596L(TTCCTC)
fusA1 G460V(GGTGTT)
fusA1 G515S(GGTAGT)
fusA1 G676S(GGTAGT)
fusA1 M461V(ATGGTG)
fusA1 N592I(AACATC)
fusA1 Q122L(CAGCTG)
fusA1 Q678L(CAGCTG)
fusA1 R59H(CGTCAT)
fusA1 R680C(CGCTGC)
fusA1 T671A(ACCGCC)
ptsP A226P(GCTCCT)
ptsP A685D(GCCGAC)
ptsP coding(1122/2280nt)
ptsP coding(1125/2295nt)
ptsP coding(12041206/2295nt)
ptsP coding(1247/2295nt)
ptsP coding(1261/2295nt)
ptsP coding(163170/2280nt)
ptsP coding(1800/2295nt)
ptsP coding(1827/2295nt)
ptsP coding(18461887/2280nt)
ptsP coding(2240/2295nt)
ptsP coding(305/2295nt)
ptsP coding(709722/2295nt)
ptsP coding(731/2295nt)
ptsP coding(847851/2295nt)
ptsP coding(889/2280nt)
ptsP G190D(GGCGAC)
ptsP G467C(GGCTGC)
ptsP G681C(GGTTGT)
ptsP P597L(CCGCTG)
ptsP T630N(ACCAAC)
ptsP V661E(GTGGAG)
ptsP Y745*(TACTAG)
cyoA G314V(GGTGTT)
ACX60_RS16660 coding(120/372nt)
PA14_58320 P110L(CCGCTG)
orfH coding(718731/1269nt)
orfH V32G(GTCGGC)
orfK coding(162/1137nt)
orfL coding(288/1113nt)
orfN coding(148/1017nt)
ACX60_RS06505 D101E(GACGAA)
ACX60_RS10565/ACX60_RS10570 intergenic(350/96)
morA E1153K(GAGAAG)
morA K1123E(AAGGAG)
PA14_50060 M1M(GTGATG)
PA14_64050 R26C(CGCTGC)
PA14_RS03500/tuf intergenic(+20/65)
PA14_RS20330/PA14_50060 intergenic(+103/14)
PA14_RS27645 Q252P(CAGCCG)
wspF E189*(GAGTAG)
Frequency
0
25
50
75
100

**Figure S2. Population sequencing reveals molecular adaptations to tobramycin and biofilm selection.** Three populations per treatment for no antibiotic lineages and five populations per treatment for antibiotic lineages were sequenced at the day twelve timepoint for each species. Shading indicates the total frequency of the mutations in each locus at day twelve.

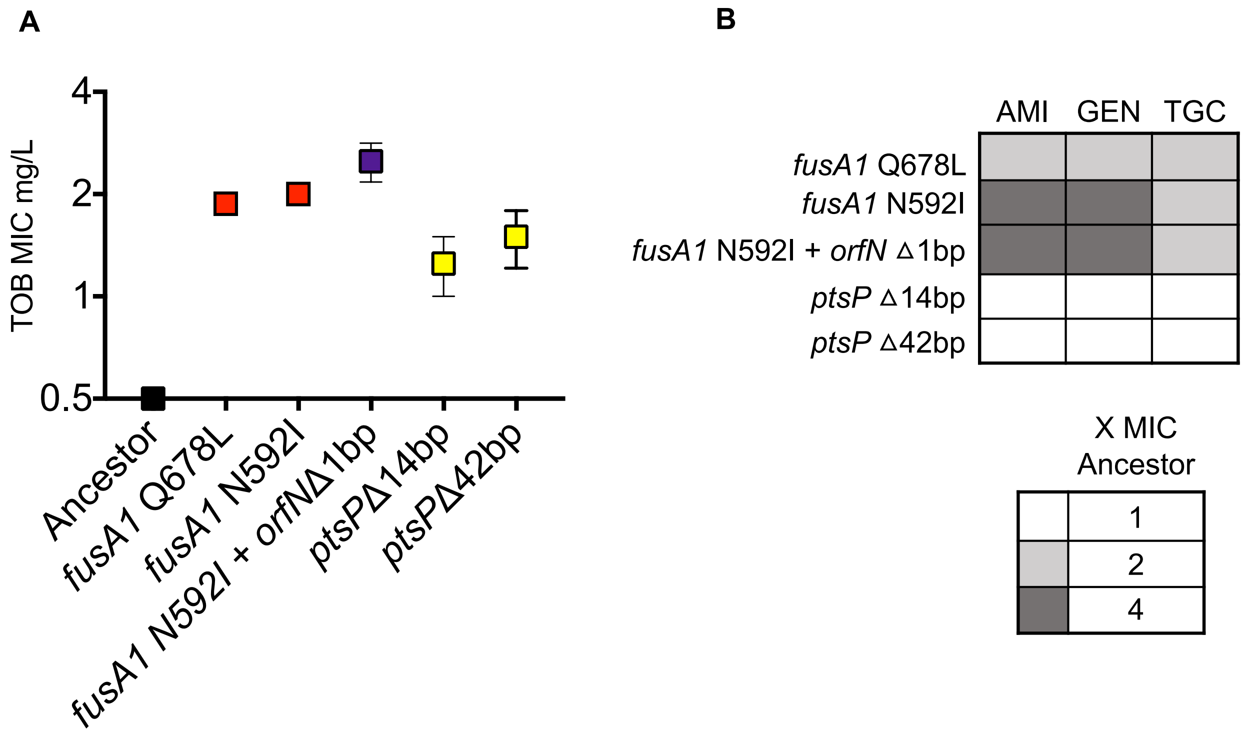


**Figure S3. MICs and relative fitness of *P. aeruginosa fusA1, fusA1+orfN,* and *ptsP* mutants compared to the ancestral genotype. A)** MICs of TOB for mutants isolated from evolved populations was determined by broth microdilution according to CLSI guidelines. The mean of at least three replicates is shown, error bars represent SEM. **B)** Change in MIC of ribosome-targeting antibiotics for genotypes selected for by TOB treatment was determined by Mueller Hinton broth microdilution using Sensititre plates. Shading indicates the average increase in MIC relative to the MIC of the ancestral clone, three replicates were performed for each genotype. *ptsP* Δ14bp corresponds to a deletion at nucleotides 1296-1309/2280, *ptsP* Δ42bp corresponds to a deletion at nucleotides 1846-1887/2280.

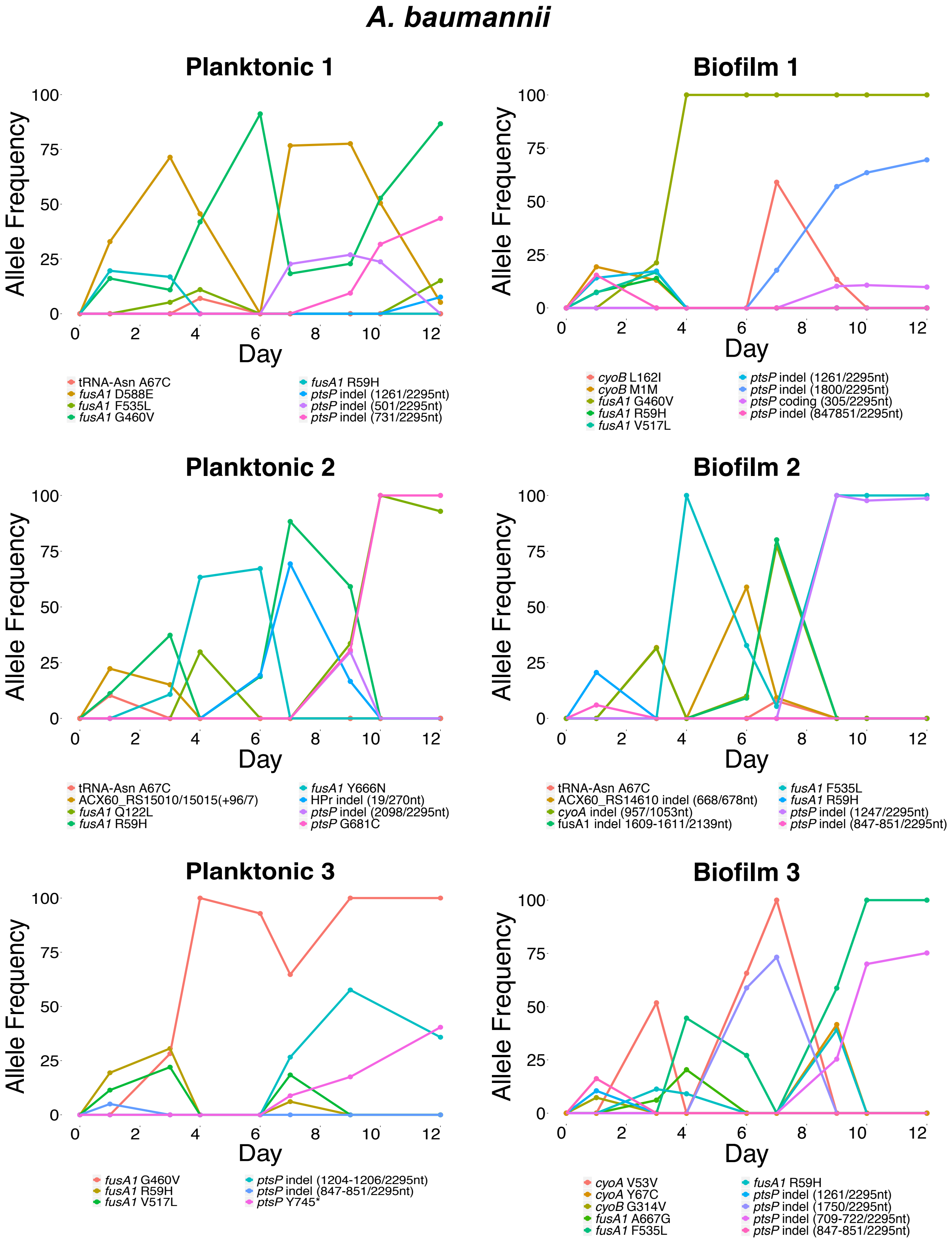
A. baumannii
Planktonic 1
Allele Frequency
Day
tRNA-Asn A67C
fusA1 D588E
fusA1 F535L
fusA1 G460V
fusA1 R59H
ptsP indel (1261/2295nt)
ptsP indel (501/2295nt)
ptsP indel (731/2295nt)
Biofilm 1
cyoB L162I
cyoB M1M
fusA1 G460V
fusA1 R59H
fusA1 V517L
ptsP indel (1261/2295nt)
ptsP indel (1800/2295nt)
ptsP coding (305/2295nt)
ptsP indel (847851/2295nt)
Planktonic 2
tRNA-Asn A67C
ACX60_RS15010/15015(+96/7)
fusA1 Q122L
fusA1 R59H
fusA1 Y666N
HPr indel (19/270nt)
ptsP indel (2098/2295nt)
ptsP G681C
Biofilm 2
tRNA-Asn A67C
ACX60_RS14610 indel (668/678nt)
cyoA indel (957/1053nt)
fusA1 indel 1609-1611/2139nt)
fusA1 F535L
fusA1 R59H
ptsP indel (1247/2295nt)
ptsP indel (847-851/2295nt)
Planktonic 3
fusA1 G460V
fusA1 R59H
fusA1 V517L
ptsP indel (1204-1206/2295nt)
ptsP indel (847-851/2295nt)
ptsP Y745*
Biofilm 3
cyoA V53V
cyoA Y67C
cyoB G314V
fusA1 A667G
fusA1 F535L
fusA1 R59H
ptsP indel (1261/2295nt)
ptsP indel (1750/2295nt)
ptsP indel (709-722/2295nt)
ptsP indel (847-851/2295nt)

### *P. aeruginosa*

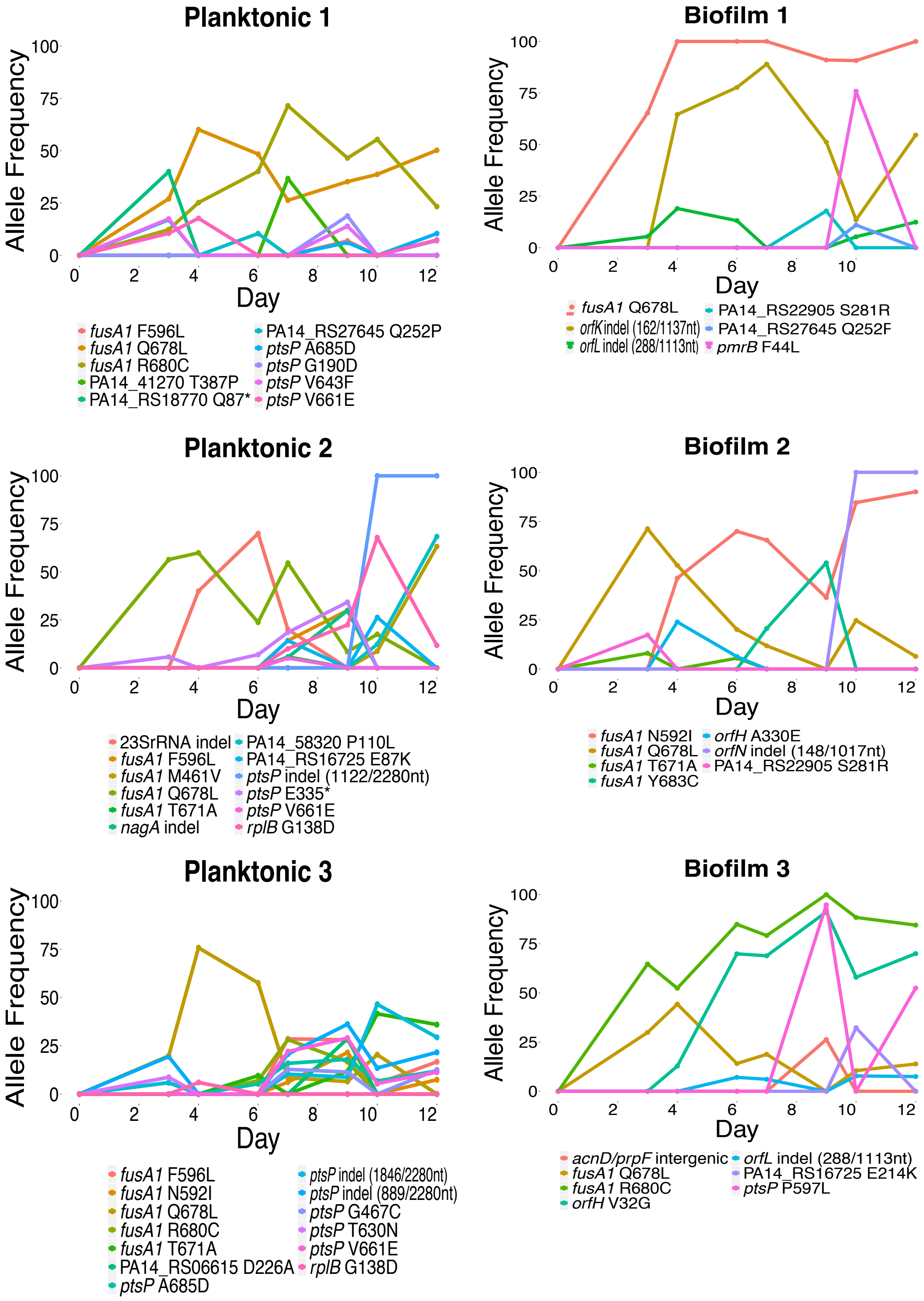

**Figure S4. Allele frequency plots of three populations from each treatment.** *A. baumannii* (top) and *P. aeruginosa* (bottom) mutations are plotted by their frequency within each population on each day of the experiment in which population sequencing was performed. SNPs are indicated by gene name followed by amino acid change and insertions or deletions are indicated by the gene name followed by "coding". In cases where insertions or deletions occurred at multiple positions within a gene, coding is followed by a number to differentiate it from other mutations insertions or deletions in that gene, for example *ptsP* coding.1 and *ptsP* coding.2.

### *fusA1*

```
A. baumannii    MARQTPITRYRNIGISAHIDAGKTTTTERILFYTGVSHKIGEVHDGAATMDWMEQEQERGITITSAATTCFWSGMGNQFPQHRINVIDTPGHVDFTIEVE
P. aeruginosa   MARTTPINRYRNIGICAHVDAGKTTTTERVLFYTGVNHKLGEVHDGAATTDWMVQEQERGITITSAAVTTFWKGSRGQYDNYRVNVIDTPGHVDFTIEVE
S. typhimurium  MARTTPIARYRNIGISAHIDAGKTTTTERILFYTGVNHKIGEVHDGAATMDWMEQEQERGITITSAATTAFWSGMAKQYEPHRINIIDTPGHVDFTIEVE
E. coli         MARTTPIARYRNIGISAHIDAGKTTTTERILFYTGVNHKIGEVHDGAATMDWMEQEQERGITITSAATTAFWSGMAKQYEPHRINIIDTPGHVDFTIEVE
S. aureus       MAREFSLEKTRNIGIMAHIDAGKTTTTERILYYTGRIHKIGETHEGASQMDWMEQEQDRGITITSAATTAAWEG-------HRVNIIDTPGHVDFTVEVE

A. baumannii    RSMRVLDGACMVYCAVGGVQPQSETVWRQANKYKVPRLAFVNKMDRTGANFFRVVEQMKTRLGANPVPIVVPIGAEDTFTGVVDLIEMKAIIWDEASQGM
P. aeruginosa   RSLRVLDGAVVVFCGTSGVEPQSETVWRQANKYGVPRIVYVNKMDRQGANFLRVVEQIKKRLGHTPVPVQLAIGAEENFVGQVDLIKMKAIYWNDDDKGM
S. typhimurium  RSMRVLDGAVMVYCAVGGVQPQSETVWRQANKYKVPRIAFVNKMDRMGANFLKVVGQIKTRLGANPVPLQLAIGAEEGFTGVVDLVKMKAINWNDADQGV
E. coli         RSMRVLDGAVMVYCAVGGVQPQSETVWRQANKYKVPRIAFVNKMDRMGANFLKVVNQIKTRLGANPVPLQLAIGAEEHFTGVVDLVKMKAINWNDADQGV
S. aureus       RSLRVLDGAVTVLDAQSGVEPQTETVWRQATTYGVPRIVFVNKMDKLGANFEYSVSTLHDRLQANAAPIQLPIGAEDEFEAIIDLVEMKCFKYTN-DLGT

A. baumannii    KFEYGEIPADLVDTAQEWRTNMVEAAAEASEELMDKYLEEGDLSKEDIIAGLRARTLASEIQVMLCGSAFKNKGVQRMLDAVIEFLPSPTEVKAIEGILD
P. aeruginosa   TYREEEIPAELKDLAEEWRSSMVEAAAEANEELMNKYLEEGELSEAEIKEGLRLRTLACEIVPAVCGSSFKNKGVPLVLDAVIDYLPAPTEIPAIKGVSP
S. typhimurium  TFEYEDIPADMQDLANEWHQNLIESAAEASEELMEKYLGGEELTEEEIKQALRQRVLNNEIILVTCGSAFKNKGVQAMLDAVIDYLPSPVDVPAINGILD
E. coli         TFEYEDIPADMVELANEWHQNLIESAAEASEELMEKYLGGEELTEAEIKGALRQRVLNNEIILVTCGSAFKNKGVQAMLDAVIDYLPSPVDVPAINGILD
S. aureus       EIEEIEIPEDHLDRAEEARASLIEAVAETSDELMEKYLGDEEISVSELKEAIRQATTNVEFYPVLCGTAFKNKGVQLMLDAVIDYLPSPLDVKPIIGHRA

A. baumannii    DKDETKASREASDEAPFSALAFKIMNDKFVGNLTFVRVYSGVLKQGDAVYNPVKSKRERIGRIVQMHANERQDIDEIRAGDIAACVGLKDVTTGDTLCDE
P. aeruginosa   DDETVEDERHADDNEPFSSLAFKIATDPFVGTLTFARVYSGVLSSGDSVLNSVKGKKERVGRMVQMHANQREEIKEVRAGDIAALIGMKDVTTGDTLCSI
S. typhimurium  DGKDTPAERHASDDEPFSALAFKIATDPFVGNLTFFRVYSGVVNSGDTVLNSVKTARERFGRIVQMHANKREEIKEVRAGDIAAAIGLKDVTTGDTLCDP
E. coli         DGKDTPAERHASDDEPFSALAFKIATDPFVGNLTFFRVYSGVVNSGDTVLNSVKAARERFGRIVQMHANKREEIKEVRAGDIAAAIGLKDVTTGDTLCDP
S. aureus       SNPEEEVIAKADDSAEFAALAFKVMTDPYVGKLTFFRVYSGTMTSGSYVKNSTKGKRERVGRLLQMHANSRQEIDTVYSGDIAAAVGLKDTGTGDTLCGE

A. baumannii    KNIITLERMEFPDPVIQLAVEPKTKADQEKMSIALGRLAKEDPSFRVHTDEESGQTIIAGMGELHLDIIVDRMKREFGVEANIGKPMVAYRETIKKTV-E
P. aeruginosa   EKPIILERMDFPEPVISVAVEPKTKADQEKMGIALGKLAQEDPSFRVKTDEESGQTIISGMGELHLDIIVDRMKREFGVEANIGKPQVAYRETITKDNVE
S. typhimurium  ENPIILERMEFPEPVISIAVEPKTKADQEKMGLALGRLAKEDPSFRVWTDEESNQTIIAGMGELHLDIIVDRMKREFNVEANVGKPQVAYREAIRAKVTD
E. coli         DAPIILERMEFPEPVISIAVEPKTKADQEKMGLALGRLAKEDPSFRVWTDEESNQTIIAGMGELHLDIIVDRMKREFNVEANVGKPQVAYRETIRQKVTD
S. aureus       KNDIILESMEFPEPVIHLSVEPKSKADQDKMTQALVKLQEEDPTFHAHTDEETGQVIIGGMGELHLDILVDRMKKEFNVECNVGAPMVSYRETFKSSA-Q

A. baumannii    QEGKFVRQTGGKGKFGHVYVRLEPLDVEAAG---KEYEFAEEVVGGVVPKEFFGAVDKGIQERMKNGVLAGYPVVGVKAVLFDGSYHDVDSDELSFKMAG
P. aeruginosa   IEGKFVRQSGGRGQFGHCWIRFSAADVDEKGNITEGLVFENEVVGGVVPKEYIPAIQKGIEEQMKNGVVAGYPLIGLKATVFDGSYHDVDSNEMAFKIAA
S. typhimurium  IEGKHAKQSGGRGQYGHVVIDMYPLEP---GSNPKGYEFINDIKGGVIPGEYIPAVDKGIQEQLKSGPLAGYPVVDLGVRLHFGSYHDVDSSELAFKLAA
E. coli         VEGKHAKQSGGRGQYGHVVIDMYPLEP---GSNPKGYEFINDIKGGVIPGEYIPAVDKGIQEQLKAGPLAGYPVVDMGIRLHFGSYHDVDSSELAFKLAA
S. aureus       VQGKFSRQSGGRGQYGDVHIEFTPNET---G---AGFEFENAIVGGVVPREYIPSVEAGLKDAMENGVLAGYPLIDVKAKLYDGSYHDVDSSEMAFKIAA

A. baumannii    SYAFRDGFMKADPVLLEPIMKVEVETPEDYMGDIMGDLNRRRGMVQGMDDLPGGTKAIKAEVPLAEMFGYATQMRSMSQGRATYSMEFAKYAETPRNVAE
P. aeruginosa   SMATKQLAQKGGGKVLEPIMKVEVVTPEDYMGDVMGDLNRRRGLIQGMEDTVSG-KVIRAEVPLGEMFGYATDVRSMSQGRASYSMEFSKYAEAPSNIVE
S. typhimurium  SIAFKEGFKKAKPVLLEPIMKVEVETPEENTGDVIGDLSRRRGMLKGQESEVTG-VKIHAEVPLSEMFGYATQLRSLTKGRASYTMEFLKYDDAPNNVAQ
E. coli         SIAFKEGFKKAKPVLLEPIMKVEVETPEENTGDVIGDLSRRRGMLKGQESEVTG-VKIHAEVPLSEMFGYATQLRSLTKGRASYTMEFLKYDEAPSNVAQ
S. aureus       SLALKEAAKKCDPVILEPMMKVTIEMPEEYMGDIMGDVTSRRGRVDGMEPRGNA-QVVNAYVPLSEMFGYATSLRSNTQGRGTYTMYFDHYAEVPKSIAE

A. baumannii    GIIAKFQAGGKKGDDE
P. aeruginosa   ALV---KKQG------
S. typhimurium  AVI---EARGK-----
E. coli         AVI---EARGK-----
S. aureus       DIIK--KNKGE-----
```

### *ptsP*

```
P. aeruginosa   ----MLNTLRKIVQEVNSAKDLKAALGIIVQRVKEAMGTQVCSVYLLDTETQRFVLMATEGLNKRSIGKVSMAPSEGLVGLVGTREEPLNLENAAAHPRY
A. baumannii    MSNMQLDTLRRIVQEINASVSLHESLDIMVNQVAEAMKVDVCSIYLLDERNQRYVLMASKGLNPESVGHVSLQLGEGLVGLVGQREEIVNLDNAPKHERF

P. aeruginosa   RYFAETGEERYASFLGAPIIHHRRVMGVLVVQQKERRQFDEGEEAFLVTMSAQLAGVIAHAEATGSIRGLGKLGKGIQEAKFVGVPGAPGVGVGKAVVVL
A. baumannii    LYLPETGEEIYNSFLGVPVMYRRKVMGVLVVQNRLPQDFSEAAESFLVTLCAQLSGVIAHAHAVGNIDVFRKPSNGPAYKTFQGVSGAGGVALGRAIILY

P. aeruginosa   PPADLEVVPDKQVDDIDAEIALFKQALEGVRADMRALSSKLASQLRKEERALFDVYLMMLDDASIGNEVKRIIRTGQWAQGALRQVVMEHVQRFELMDDA
A. baumannii    PPADLGSVPDREAEDISDELRILDQAISSVRSEIRSLDEKMHDSLMAEERALFSVFLRMLDENALPAEIKELIRDGHWAQGAVRRVIEKHTALFAQMEDD

P. aeruginosa   YLRERASDVKDIGRRLLAYLQEERKQNLTYPEQTIIVSEELSPAMLGEVPEGRLVGLVSVLGSGNSHVAILARAMGIPTVMGAVDLPYSKVDGIDLIVDG
A. baumannii    YLRERVSDLKDLGRRILAYLQEEDSSHRELSPDSILIGEEISTAALVELPVDNIAAIVTSEGAANSHMVIVARALGIPTVVGVTELPVNTLDDAEMIVDA

P. aeruginosa   YHGEVYTNPSAELVRQYSDVVAEERELSKGLAALRELPCETLDGHRMPLWVNTGLLADVARAQERGAEGVGLYRTEVPFMINDRFPSEKEQLAIYREQLS
A. baumannii    YQGRVFVNPPRRLRQRYKEIQKEDEQIAKDLKQYETKEAITPDGVSVQLFVNTGLMIDVVRGVQRGAQGVGLYRSEIPFMLRERFPGEEEQRAIYRQQLS

P. aeruginosa   AFHPLPVTMRTLDIGGDKALSYFPIKEDNPFLGWRGIRVTLDHPEIFLVQTRAMLKASEGLDNLRILLPMISGTHELEEALHLIHRAWGEVRDE-GVDIA
A. baumannii    HFANKPVVMRTLDIGADKDLPYFSIEEENSALGWRGIRFTLDHPEIFSAQIRAMLKASIGLNNLHILLPMVTTVSEVEEVLYLLERDWIAVQEEEQVKIT
                6
P. aeruginosa   MPPIGMMVEIPAAVYQTRELARQVDFLSVGSNDLTQYLLAVDRNNPRVADLYDYLHPAVLHALKKVVDDAHLEGKPVSICGEMAGDPAAAVLLMAMGFDS
A. baumannii    KPKIGIMVEVPSVLLQIDEFAELVDFFSVGSNDLTQYLLAVDRNNPHVANVYSHFHPSILRALTRLVKECHKYQKPVSICGEMAGDPLSAILLMAMGFNT

P. aeruginosa   LSMNATNLPKVKWLLRQITLDKARDLLGQLLTFDNPQVIHSSLHLALRNLGLGRVINPAATVQP
A. baumannii    LSMSSSNILRVRKAICHVPMSDAQKLLDDVMKMNNPLIVKSWLEYYFKTHGLADMVKSNRLVSV
```

### *cyoA*

```
A. baumannii    ----MRQTILAVLSLSTLAALLTGCGGDMVLLNSKGPVGQGQSDLMMTAIYLMLLVVIPSIIMALWFGWKYRASNKDADYKPTWAHSTAIEVVWGIPVI
P. aeruginosa   MTKANPFAALKWLSLAP-ALLLGGC--DMTLFNPKGQVGMDERTLIITATLLMLIVVIPVIVMTLAFAWKYRASNTQAEYKPDWHHSNRIEAVWLVPCV
S. enterica     --------------------MLSGC--NSALLDPKGQIGLEQRSLILTAFGLMLIVVIPAILMAVGFAWKYRASNKDAKYSPNWSHSNKVEAVWTVPIL
E. coli         ---------MGWLSLIAGTALLSGC--NSALLDPKGQIGLEQRSLILTAFGLMLIVVIPAILMAVGFAWKYRASNKDAKYSPNWSHSNKVEAVWTVPIL

A. baumannii    IIGILAWLTWWGSHKYDPYRPLESDKAPLTIQVIAEQFKWIFIYPEQNIATVNEVRFPEKTPLSFKITSNFTMNSFFIPQLGGQIYAMAGMQTHLHLLAN
P. aeruginosa   IIAILGWITWESTHKLDPYRPLDSEVKPVTIQAVSLDWKWLFIYPEQGIATVNEIAFPKDTPVNFQITSDSVMNSFFIPQLGSQIYSMAGMMTKLHLIAN
S. enterica     IIIFLAVLTWKTTHALEPSKPLAHDEKPITIEVVSMDWKWFFIYPEQGIATVNEIAFPANTPVYFKVTSNSVMNSFFIPRLGSQIYAMAGMQTRLHLIAN
E. coli         IIIFLAVLTWKTTHALEPSKPLAHDEKPITIEVVSMDWKWFFIYPEQGIATVNEIAFPANTPVYFKVTSNSVMNSFFIPRLGSQIYAMAGMQTRLHLIAN

A. baumannii    ETGVYRGFSSNYSGYGFSQMRFKAHSVTE-QQFNEWVAAVKAGNGTTINPEAVQKTTLDQAELATLRDGDRSKHQIEHLVNRAKAAGDQEALAKAEAMKP
P. aeruginosa   EEGVFDGISANYSGGGFSGMRFKAIATSE-QGFQDWVAKVKAAPTSLS-IGTYPELVKPSENVPPTYFSSVSPELFGHILTKYEHHGDAKGAAHGEHAG-
S. enterica     EPGTYDGISASYSGPGFSGMKFKAIATPDRAAFDQWVAKAKQSPNTMSDMAAFEKLAAPSEYNQVEYFSNVKPDLFADVINKFMAHGKSMDMTQPEGEH-
E. coli         EPGTYDGISASYSGPGFSGMKFKAIATKDRAEFDQWVAKAKQSPNTMNDMAAFEKVAMPSEYNKVEYFSSVKPDLFKDVINKFMGHGKSMDMTQPEGEH-

A. baumannii    FPTKPHPVTYYSSVEPKLFETIINHYMSNYHGADHSAAHTAAETHVAAEHAAQGE
P. aeruginosa   -----------AEHEAAMTG-------HDMQDMDMQAMQGMKDMKDMHMQPSTQE
S. enterica     -----------SAHEG-------------MEGMDMSHAESAH-------------
E. coli         -----------SSHEG-------------MEGMDMSHAESANSKG----------
```

***cyoB***

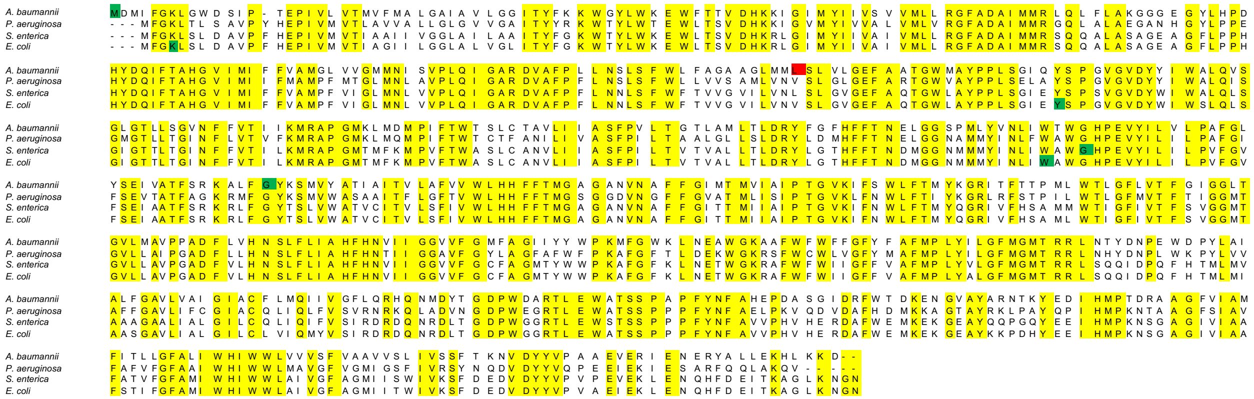


**Figure S5. Multiple sequence alignment of *fusA1*, *ptsP, cyoA*, and *cyoB* for the species in which mutations have been demonstrated to confer aminoglycoside resistance.** Positions with identical amino acid identities across all species are highlighted in yellow. SNPs identified in this experiment, previous *in vitro* experiments, and clinical isolates are shown at their relative positions within the amino acid sequence. SNPs in conserved positions are highlighted in green and SNPs in non-conserved positions are highlighted in red. SNPs occur in conserved positions more frequently than expected based on the frequency of conserved positions in *fusA1*, *ptsP*, and *cyoA*: *fusA1* ($\chi^2$ = 11.58, df = 1, p = 0.0006), *ptsP* ($\chi^2$ = 4.37, df = 1, p = 0.03652), *cyoA* ($\chi^2$ = 3.89, df = 1, p = 0.0486), *cyoB* ($\chi^2$ = 2.53, df = 1, p = 0.1114).

| **Genome Name** | **PATRIC Genome ID** | **NCBI Reference Genome ID** | **Gene** | **Product** | **Old RefSeq Locus Tag** | **New RefSeq Locus Tag** | **Start** | **End** | **NT Length** | **Strand** | **AA Length** | **Non-redundant protein accession number** |
|---|---|---|---|---|---|---|---|---|---|---|---|---|
| *Pseudomonas aeruginosa* UCBPP-PA14 | 208963.12 | NC_008463 | *fusA1* | Translation elongation factor G | PA14_08820 | PA14_RS03565 | 755665 | 757785 | 2121 | + | 706 | WP_003093741 |
| | | | *ptsP* | Phosphocarrier protein kinase/phosphorylase, nitrogen regulation associated | PA14_04410 | PA14_RS01795 | 392770 | 395049 | 2280 | + | 759 | WP_003084404 |
| | | | cyoB | Cytochrome O ubiquinol oxidase subunit I | PA14_47190 | PA14_RS19185 | 4203575 | 4205551 | 1977 | - | 658 | WP_003082705 |
| | | | cyoA | Cytochrome O ubiquinol oxidase subunit II | PA14_47210 | PA14_RS19190 | 4205558 | 4206553 | 996 | - | 331 | WP_003086856 |
| *Acinetobacter baumannii* strain ATCC 17978-mff | 470.1575 | CP012004 | *fusA* | Translation elongation factor G | ACX60_14045 | ACX60_RS14040 | 2962952 | 2965090 | 2139 | - | 712 | WP_005229047 |
| | | | *ptsP* | Phosphocarrier protein kinase/phosphorylase, nitrogen regulation associated | ACX60_16055 | ACX60_RS16050 | 3393072 | 3395366 | 2295 | + | 764 | WP_005133238 |
| | | | cyoB | Cytochrome O ubiquinol oxidase subunit I | ACX60_06730 | ACX60_RS06730 | 1439111 | 1441102 | 1992 | - | 663 | WP_002119744 |
| | | | cyoA | Cytochrome O ubiquinol oxidase subunit II | ACX60_06735 | ACX60_RS06735 | 1441106 | 1442158 | 1053 | - | 350 | WP_005215377 |
| *Escherichia coli* str. K-12 substr. MG1655 | 511145.12 | NC_000913 | *fusA* | Translation elongation factor G | b3340 | | 3469422 | 3471536 | 2115 | - | 704 | WP_000124700 |
| | | | cyoB | Cytochrome O ubiquinol oxidase subunit I | b0431 | | 447874 | 449865 | 1992 | - | 663 | WP_000467180 |
| | | | cyoA | Cytochrome O ubiquinol oxidase subunit II | b0432 | | 449887 | 450774 | 888 | - | 295 | WP_001239436 |
| *Salmonella enterica* serovar Typhimurium str. LT2 | 99287.12 | NC_003197 | *fusA* | Translation elongation factor G | STM3446 | | 3599558 | 3601672 | 2115 | - | 704 | WP_000124693 |
| | | | cyoB | Cytochrome O ubiquinol oxidase subunit I | STM0442 | | 495189 | 497180 | 1992 | - | 663 | WP_000467158 |
| | | | cyoA | Cytochrome O ubiquinol oxidase subunit II | STM0443 | | 497191 | 498120 | 930 | - | 309 | WP_001239449 |
| *Staphylococcus aureus* NCTC 8325 strain | 93061.132 | LS483365 | *fusA* | Translation elongation factor G | NCTC8325_00491 | | 540862 | 542943 | 2082 | + | 693 | WP_001788222 |

**Table S1.** Genome name and NCBI ID's for genomes referenced in this study. Accession numbers and locus tags for driver mutations of aminoglycoside resistance identified in this study for each of these species are indicated. Nucleotide length, amino acid length, position within the genome, and strand are also shown.
